# Cadherin-6 mediates thrombosis *in vivo*

**DOI:** 10.1101/2020.04.20.051045

**Authors:** Emma G. Bouck, Maria de la Fuente, Elizabeth R. Zunica, Wei Li, Michele M. Mumaw, Marvin T. Nieman

**Affiliations:** Department of Pharmacology, Case Western Reserve University, Cleveland, United States; Deparmtent of Biomedical Sciences, Marshall University Joan C. Edwards School of Medicine, Huntington, West Virginia, United States

**Keywords:** cadherin-6, arterial thrombosis, platelet adhesion, mouse model

## Abstract

**Background:** Platelet adhesion is the critical process mediating stable thrombus formation. Previous reports of cadherin-6 on human platelets have demonstrated its role in platelet aggregation and thrombus formation.

**Objectives:** We aimed to further characterize the importance of cadherin-6 in thrombosis *in vivo*.

**Methods:** Cadherin-6 platelet expression was evaluated by western blotting, flow cytometry and immunoprecipitation. Thrombosis was evaluated using the FeCl_3_ and Rose Bengal carotid artery models in C57Bl6 mice treated with anti-cadherin-6 or IgG and wild-type or *Cdh6^-/-^* mice. Platelet function was compared in wild type and *Cdh6^-/-^* mice using tail-clip assays and aggregometry.

**Results:** Human platelet expression of cadherin-6 was confirmed at ~3,000 copies per platelet. *Cdh6^-/-^* mice or those treated with anti-Cadherin-6 antibody showed an increased time to occlusion in both thrombosis models. Cadherin-6 was not expressed on mouse platelets, and there were no differences in tail bleeding times or platelet aggregation in wild-type versus *Cdh6^-/-^* mice.

**Conclusions:** Cadherin-6 plays an essential role in thrombosis *in vivo*. However, cadherin-6 is not expressed on murine platelets. These data are in contrast to human platelets, which express a functional cadherin-6/catenin complex. The essential, platelet-independent role for cadherin-6 in hemostasis may allow it to be an effective and safe therapeutic target.

**Essentials:**

- Cadherin-6 function in thrombus formation was investigated in vivo using two murine models of thrombosis.
- Blocking or deleting cadherin-6 significantly delayed time to occlusion
- Human platelets express cadherin-6, but murine platelets do not.

## 1. Introduction

Arterial thrombosis occurs following rupture of an atherosclerotic plaque. Platelets are activated upon contact with the damaged vessel and potentiate thrombus formation. A primary consequence of platelet activation is the conformational change of platelet integrins to a high-affinity state.[1] The most essential and abundant platelet integrin is α_llb_β_3_, which builds platelet aggregates by crosslinking fibrinogen.[2]

Cadherin-6 was proposed as a novel platelet-expressed ligand to α_llb_β_3_.[3] Junctional proteins, including cadherins, were first documented on platelets by Elrod et al.[4] E-cadherin has also been identified on platelets.[5] Cadherins are calcium-dependent, single-pass transmembrane adhesion proteins with characteristic extracellular repeats (ECs).[6] Cadherin-6 is a type II classical cadherin, containing 5 tandem ECs and two tryptophan residues in EC1 which facilitate trans-cadherin binding.[7] Notably, cadherin-6 contains an Arg-Gly-Asp (RGD) tripeptide in EC1.[8] RGD domains are well-characterized integrin binding motifs; α_llb_β_3_ binds to RGD sequences on fibrinogen.[9]

A supportive role for platelet cadherin-6 in platelet aggregation and thrombus formation was demonstrated by platelet adhesion to immobilized cadherin-6.[3] The current understanding of cadherin-6 in platelet adhesion is solely based on *in vitro* and *ex vivo* studies. The goal of this study was to examine the role of cadherin-6 *in vivo*. Herein, we demonstrate that cadherin-6 has a significant role in thrombus formation *in vivo*. However, analysis of murine platelets exposed an absence of cadherin-6, revealing an important species difference and a platelet-independent role for cadherin-6 in thrombosis.

## 2. Methods

### 2.1 Animals

C57BL/6J and Cdh6^-/-^ (B6.129S6-Cdh6^tm1sma^) mice were purchased from The Jackson Laboratory (Bar Harbor, Maine). Platelets from FVB and 129×1/SVJ mice were also used to assess cadherin-6 expression. All animal studies were approved by the Institutional Animal Care and Use Committee at Case Western Reserve University School of Medicine.

### 2.2 Preparation of platelet-rich plasma (PRP)

With approval by the Case Western Reserve University Institutional Review Board, human whole blood was obtained by venipuncture from healthy donors after obtaining informed consent. Murine whole blood was collected retro-orbitally; platelet count was obtained using a Hemavet 950FS (Drew Scientific Inc.). Whole blood was collected into either acid citrate dextrose (ACD) (2.5% sodium citrate, 71.4 mM citric acid, 2% D-glucose) or sodium citrate (0.15M). The blood was centrifuged at 200 x *g* for 15 min (human) or 2300 x *g* for 20 seconds (murine) at room temperature to obtain PRP. Platelet concentrations were quantified using a Coulter Counter.

### 2.3 Western blot

5×10^7^ platelets in PRP were directly suspended into SDS Laemmli buffer. CHO cells were transfected with 5 ug of human cadherin-6 in pIRES-puro or mouse cadherin-6 in pCDNA3.1 via Lipofectamine 2000 (Invitrogen). After 48hrs, the cells were lysed in RIPA buffer (1% NP-40, 0.5% Deoxycholate, 0.1% SDS) and 100 ug of protein was loaded.

For quantitative Western blotting, recombinant human Cadherin 6 Fc chimera protein (~130 kD) or recombinant mouse Cadherin 6 protein (~90 kD) was used to generate a standard curve (R&D Systems catalog # 2715-CA or 6826-CA, respectively). For mouse samples, the gel was loaded with 0.0023, 0.0115, 0.023, 0.046, and 0.23 μg of protein per lane. Based on loading 5×10^7^ platelets, this is the equivalent of 300, 1 500, 3 000, 6 000, or 30 000 molecules/platelet respectively. The same strategy was used for human samples. The formula is shown below.

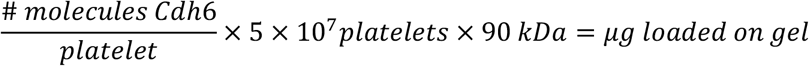

Samples were resolved by SDS-PAGE, transferred to nitrocellulose, probed with mouse monoclonal anti-cadherin-6 antibody (1:500, R&D Systems, catalog # MAB2715), then IRDYE 800CW donkey anti-mouse IgG (LI-COR Biosciences) and imaged using the Odyssey Infrared Imaging System.

Human platelet lysate was immunoprecipitated using IgG or anti-cadherin-6 antibody (R&D Systems, catalog # MAB2715) and Protein A beads (Invitrogen). The immunocomplexes were washed 3 times with Tris buffered saline with 0.1% Triton X-100 and resuspended in 2x sample buffer, and resolved on SDS-PAGE. The membrane was probed with anti-cadherin-6, anti-α-catenin (Life Technologies, catalog # MAS-161986), and anti-β-catenin (Cell Signaling, catalog #9582) antibodies.

### 2.4 Flow cytometry

Platelet rich plasma (PRP) was stimulated with SFLLRN (25 uM), AYPGKF (250 uM), convulxin (5 nM), or ADP (5 uM). Platelets were incubated with APC-conjugated anti-cadherin-6 antibody (R&D systems, catalog # FAB2175A) and analyzed using a BD LSRFortessa^™^. Cadherin-6 expression was quantified using Quantum Simply Cellular beads (Bangs Laboratories, Inc) to generate a standard curve of antibody binding sites as previously described.[10–12]

### 2.5 Carotid artery thrombosis models

Mice (8-12 weeks) were anesthetized with ketamine (100 mg/kg) and xylazine (10 mg/kg) and the carotid artery was exposed. In some experiments, mice received sheep polyclonal anti-cadherin-6 (R&D, catalog # AF2715) or sheep IgG intravenously 15 minutes prior to initiation of thrombosis. Thrombosis was induced via the FeCl_3_ model[13,14] with a minute-long topical application of filter paper saturated with 7.5% FeCl_3_ and vessel occlusion was monitored for 30 minutes via intravital microscopy. Images were captured with a QImaging Retigo Exi 12-bit mono digital camera and Streampix version 3.17.2 (Norpix).

In the Rose Bengal model[15] 4,5,6,7-tetrachloro-3, 6-dihydroxy-2,4,5,7-tetraio-dospiro (isobenzofuran-1(3H),9[9H] xanthan)-3:1 dipotassium salt (50 mg/kg in 0.9% saline, Fisher Scientific) was injected retro-orbitally before catalyzing vessel injury with a 540 nm laser. A doppler flow probe was used to monitor vessel occlusion for 90 minutes.

Because some animals did not reach full occlusion (right-censored data), the data were plotted on Kaplan-Meier survival curves and the Log-rank (Mantel-Cox) Test was used to compare curves.[16]

#### Tail-clip assay

3 mm was clipped from the tails of anesthetized wild-type and *Cdh6^-/-^* mice, and the tail was placed in 37° saline. Time to cessation of bleeding was observed for 10 minutes.

#### Platelet aggregometry

PRP was stimulated with AYPGKF (50 uM) or convulxin (0.5 nM). Platelet aggregation was measured under constant stirring (1200 rpm) with a Chrono-log Model 700 aggregometer using Aggrolink8 version 1.3.98.

## 3. Results and Discussion

### 3.1 Cadherin-6 is present on human platelets

Previous reports have observed cadherin-6 expression on human platelets.[3] To examine cadherin-6 on human platelets, we designed a western blot to quantify the density of cadherin-6 expression. Purified recombinant human cadherin-6 was used to generate a standard curve to determine the threshold of detection. Cadherin-6 was detected in CHO cells transiently transfected with human cadherin-6 and human platelets isolated from healthy donors (Figure 1A). Based on the standard curve, cadherin-6 expression is ~3000 copies per platelet. Platelet surface expression of cadherin-6 was analyzed by quantitative flow cytometry using an APC-conjugated antibody and confirmed the density of cadherin-6 to be ~3,000 copies per platelet (Figure 1B). Immunoprecipitation of cadherin-6 confirmed associations with α-catenin and β-catenin (Figure 1C). Cadherin-catenin adhesion complexes can link external stimuli to actin cytoskeleton dynamics.[17] Together, these results support functional expression of cadherin-6 on human platelets.

**Figure 1.**
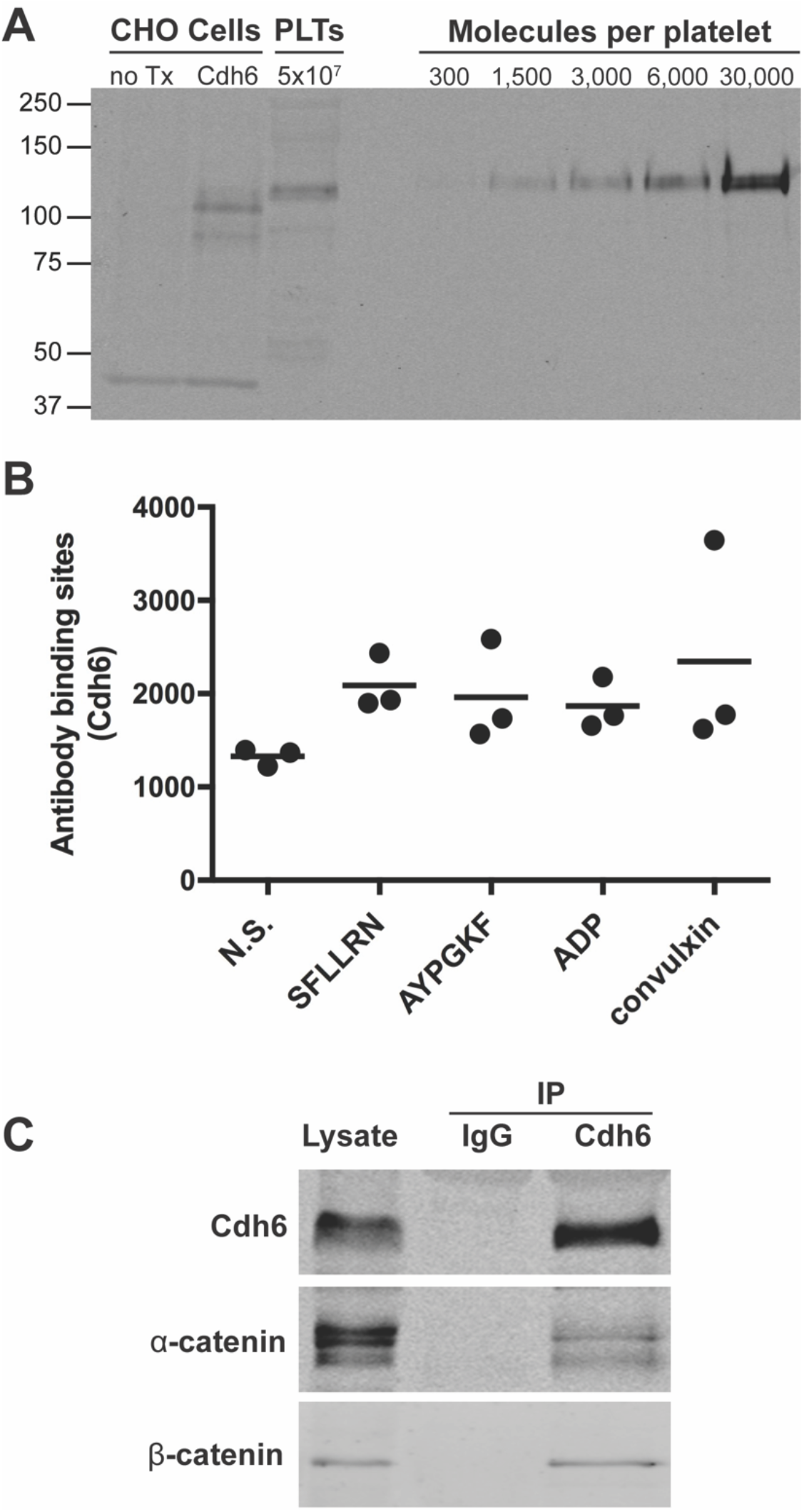
Human platelets express cadherin-6. (A) Human platelet lysate was probed with mouse monoclonal anti-cadherin-6, controlled with CHO cells transfected with human cadherin-6 (Cdh6) and recombinant human cadherin-6 Fc chimera protein. (B) Quantitative flow cytometry was performed using an APC-conjugated anti-cadherin-6 antibody on non-stimulated platelets or those activated with PAR activating peptides, ADP, or collagen. (C) Human platelet lysate was immunoprecipitated with anti-cadherin-6 antibody. Cadherin 6, α-catenin, and β-catenin were detected by western blot.

### 3.2 *Cadherin-6 has an essential role during thrombosis* in vivo

Previously, fibrinogen- and von Willebrand factor-deficient mice were treated with sheep polyclonal anti-cadherin-6 and evaluated *ex vivo* for thrombus size. Compared to the IgG control treatment, blocking cadherin-6 resulted in smaller thrombi.[3] To define the effect of blocking cadherin-6 *in vivo*, wild-type mice were treated with sheep polyclonal anti-cadherin-6 antibody or sheep IgG. Thrombosis was induced in the carotid artery using the FeCl_3_ or Rose Bengal model. In the FeCl_3_ model, mice treated with anti-cadherin-6 showed impaired accumulation of platelets at the injured vessel (Fig 2A) and 66% failed to form a stable occlusion after 30 minutes (Fig 2B). Similarly, 75% of anti-cadherin-6-treated mice never formed a full occlusion in the Rose Bengal model (Fig 2C).

**Figure 2.**
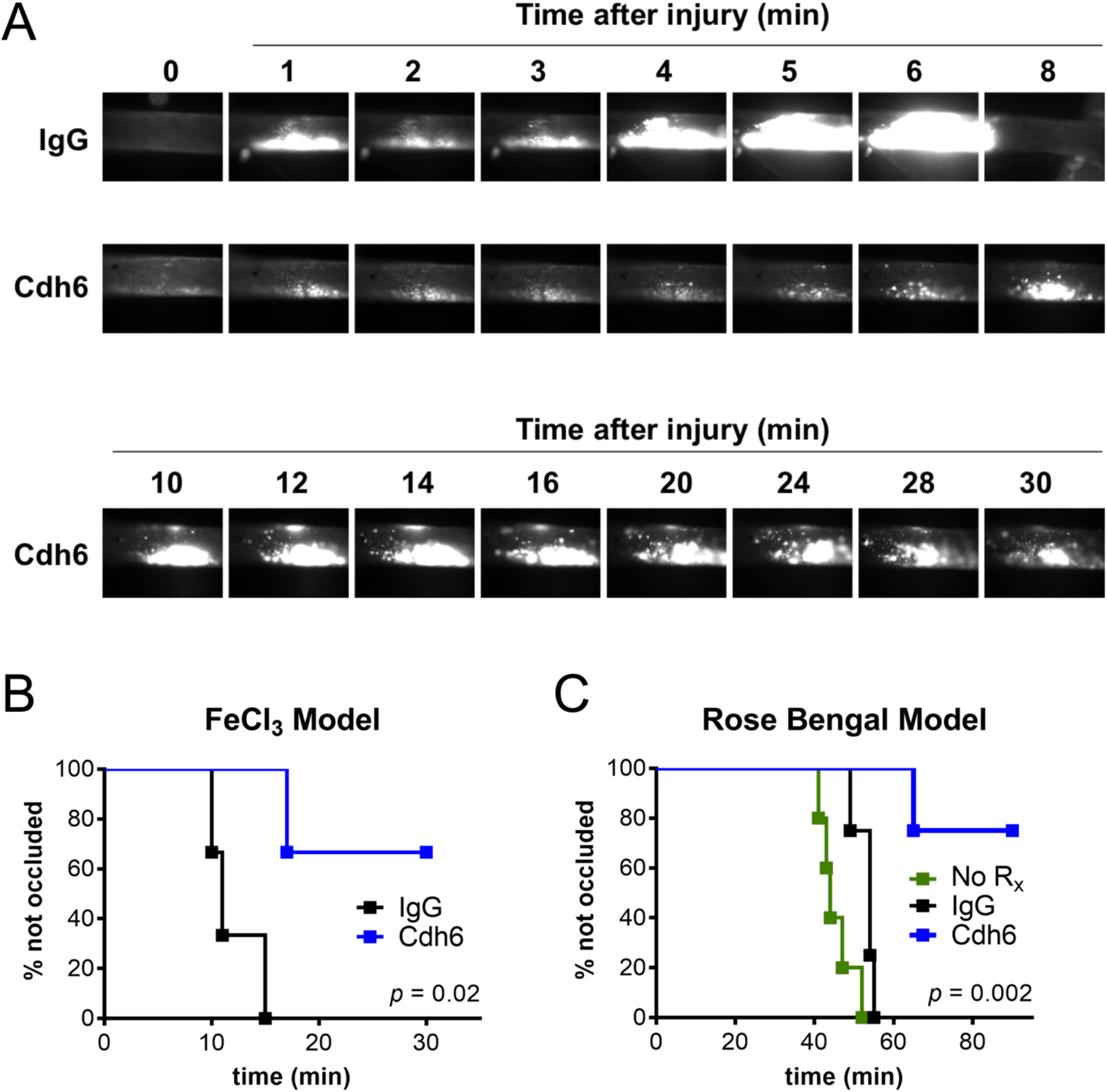
Blocking cadherin 6 disrupts thrombosis. C57BL/6J mice were treated with sheep polyclonal anti-cadherin-6 (Cdh6) 15 min prior to initiating thrombosis with FeCl_3_ or Rose Bengal. (A) Intravital microscope images obtained following FeCl_3_ treatment. Time to stable vessel occlusion was observed and represented as Kaplan-Meier curves for (B) FeCl_3_ (n=3 per group) and (C) Rose Bengal models (n=5 for no treatment, n=4 for IgG or Cdh6). The curves were compared using the Mantel-Cox Log-rank test.

To further explore the significance of cadherin-6 *in vivo*, we evaluated thrombosis in global cadherin-6 knockout *(Cdh6^-/-^)* mice. In the FeCl_3_ model, 55% of *Cdh6^-/-^* mice failed to form a stable occlusion after 30 minutes, whereas wild-types reached full occlusion at 10±2 minutes (Fig 3A). In the Rose Bengal model, loss of cadherin-6 resulted in significantly delayed occlusion; wild-types formed full occlusions in 35±2.5 min whereas *Cdh6^-/-^* mice formed full occlusions in 51.8±4.7 min (*p*=0.03) (Fig 3B). Interestingly, antibody treatment caused a greater defect in thrombosis than genetic deletion of cadherin-6, which may be explained by off-target effects of the antibody. Nonetheless, thrombosis was impaired in both models when cadherin-6 was knocked out globally, indicating a significant role of cadherin-6 *in vivo*.

**Figure 3.**
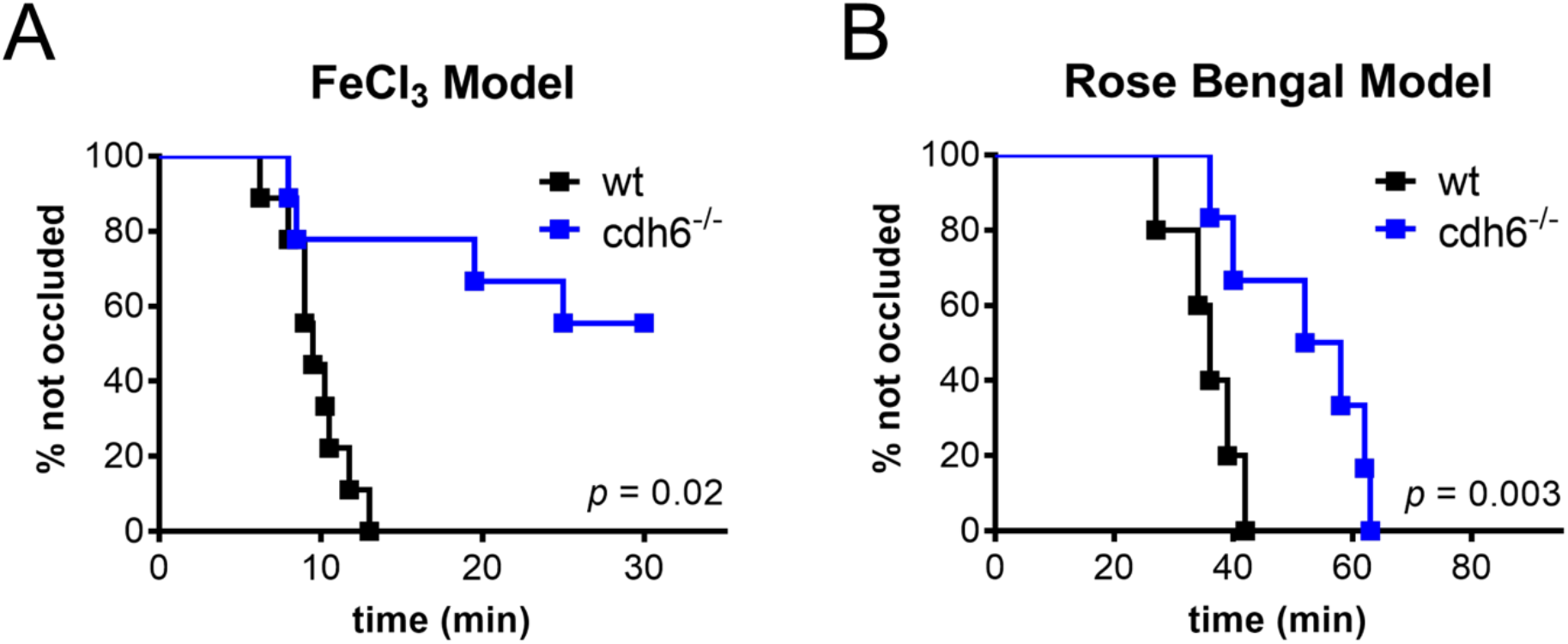
Thrombosis is delayed in CDH6 knockout mice. Thrombosis was initiated in *Cdh6^-/-^* and wild-type mice using the (A) FeCl_3_ (n=9 per group) or (B) Rose Bengal (n=5 for wild-type and n=6 for *Cdh6^-/-^*) models and time to vessel occlusion was observed. The vessels were monitored until a full occlusion was formed or the experiment ended and represented as Kaplan-Meier curves. The curves were compared using the Mantel-Cox Log-rank test.

### 3.3 Cadherin-6 is not present on murine platelets

Given our observations that cadherin-6 is expressed on human platelets and contributes to thrombosis in mice, we examined cadherin-6 expression and function on murine platelets. A western blot was performed using CHO cells transfected with murine cadherin-6, platelets isolated from wild-type and *Cdh6^-/-^* mice, and purified recombinant murine cadherin-6. Surprisingly, there was no expression of cadherin-6 in platelets from wild-type C57BL/6J mice (Fig 4A). The standard curve generated by the recombinant protein demonstrated that sensitivity of the western blot is at least 300 of copies per platelet. Western blotting additionally confirmed the absence of cadherin-6 expression on platelets from FVB and 129×1/SVJ mice (data not shown).

**Figure 4.**
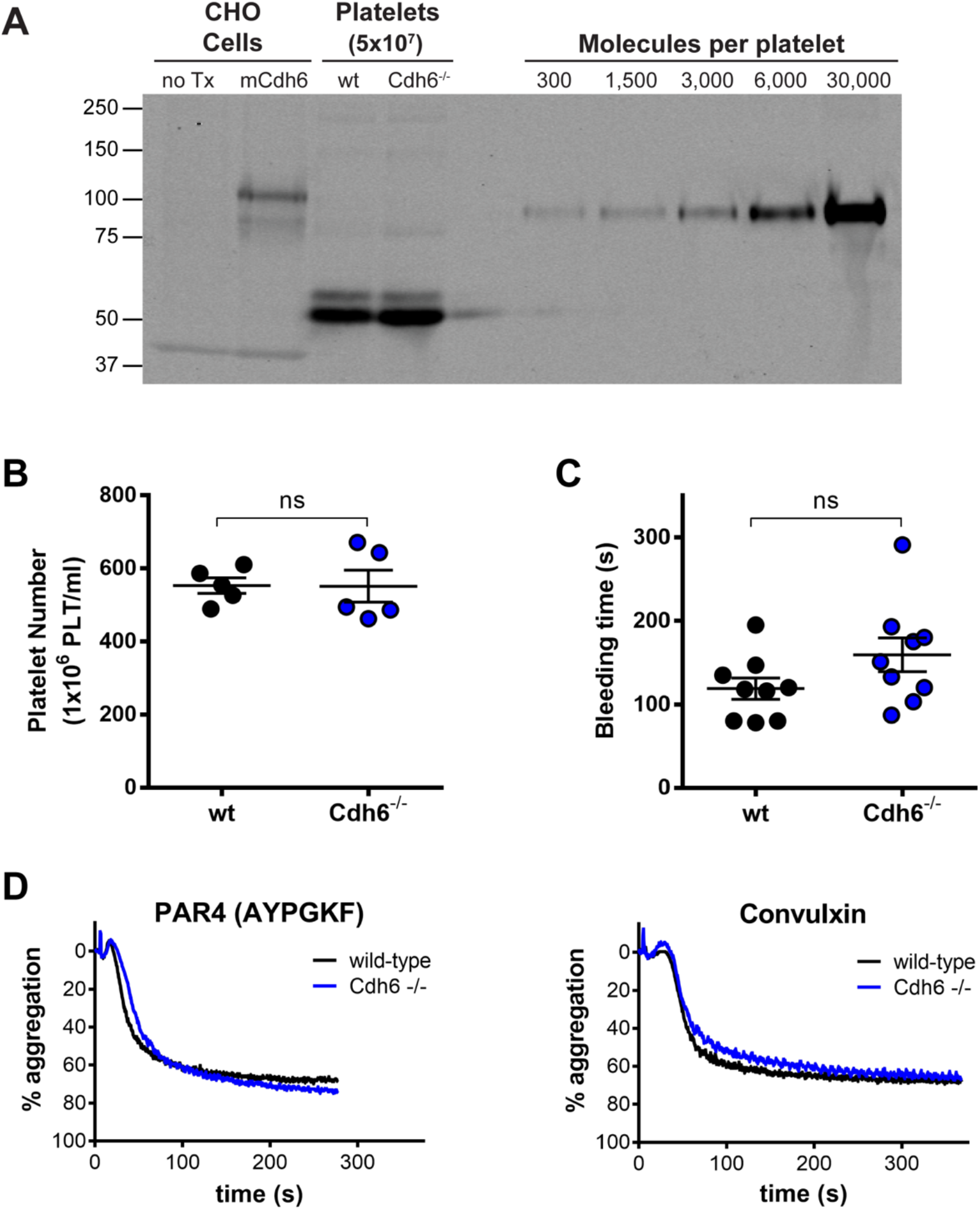
Cadherin 6 is absent on murine platelets. (A) Platelet lysates from *Cdh6^-/-^* and wildtype mice were probed with mouse monoclonal anti-cadherin-6, controlled with CHO cells transfected with mouse cadherin-6 (Cdh6) and recombinant mouse cadherin-6 protein. (B) Platelet count was measured in whole blood from wild-type and *Cdh6^-/-^* mice. (C) *Cdh6^-/-^* and wild-type mice were anesthetized before clipping 3 mm of tail to measure time to bleeding cessation. (D) *Cdh6^-/-^* and wild-type platelets were stimulated with PAR4 agonist peptide (AYPGKF) or convulxin, and platelet aggregation was measured.

The absence of cadherin-6 on platelets implies that wild-type and *Cdh6^-/-^* platelets should be indistinguishable in platelet function assays. There was no difference in platelet count between wild-type and knockout mice (Fig 4B). There was not a significant difference in the bleeding time yielded by the tail-clip assay (Fig 4C). Platelet aggregation induced with either PAR4-activating peptide or convulxin was identical in wild-type and *Cdh6^-/-^* mice (Fig 4D). Together, these results confirm cadherin-6 is not expressed on mouse platelets.

### 3.4 Discussion

Although cadherin-6 expression differs on human and murine platelets, platelets from both species adhered to immobilized cadherin-6.[3] Our results support a role for cadherin-6 in platelet adhesion. Together, significantly delayed occlusion in *Cdh6^-/-^* mice and absence of cadherin-6 on murine platelets indicate cadherin-6 expression on a cell type that supports platelet adhesion to the injured vessel. Cadherin-6 has been identified in vascular smooth muscle cells[18], and Notch3 signaling was shown to control cadherin-6 vessel expression.[19] Identifying the cell type responsible for cadherin-6-dependent platelet adhesion in murine thrombosis will lend further insight into mechanisms of cadherin-6 in human thrombosis.

Cadherin-6 is likely participating in the robust network of adhesion interactions that stabilize a developing thrombus. Since cadherin-6 contains an RGD integrin binding motif in EC1, it likely interacts heterotypically with platelet α_llb_β_3_. Given human platelet cadherin-6 expression, homotypic adhesion may reinforce and stabilize platelet aggregates at the vessel wall. Further defining these interactions will help to evaluate the potential therapeutic utility of cadherin-6.

Platelet-cadherin-6 interactions may be important in settings beyond arterial thrombosis. Tumor cells can activate platelets to promote tumor growth, invasion, and immune evasion.[20] Cadherin-6 overexpression has been identified in ovarian and renal cancers[21,22], inciting the possibility that tumor-expressed cadherin-6 mediates tumor-platelet crosstalk. Treatment of colorectal cancer or melanoma cells with an antibody targeting the cadherin-17 RGD motif reduced their metastatic potential *in vivo*.[23] Inhibiting tumor-platelet crosstalk is a promising strategy to limit tumor progression and metastasis; RGD cadherins may be promising molecular targets.

Herein, we have demonstrated an essential role for cadherin-6 for thrombosis *in vivo*. Cadherin-6 remains a poorly understood member of the cadherin superfamily, although it carries significant therapeutic potential in multiple clinical settings. Understanding the expression, regulation, and participation of cadherin-6 across different pathologies is necessary for harnessing its potential.

## Acknowledgements

This study was supported by the NIH-NHLBI grant HL098217 to MTN.

## Author contributions

EGB, WL, and MTN designed the experiments. EGB, MF, WL, ERZ, MMM, and MTN conducted experiments and analyzed the data. EGB and MTN wrote the manuscript. MF, WL, ERZ, and MMM provided critical feedback on the manuscript.

## References

1 Xu XR, Carrim N, Neves MAD, McKeown T, Stratton TW, Coelho RMP, Lei X, Chen P, Xu J, Dai X, Li BX, Ni H. Platelets and platelet adhesion molecules: Novel mechanisms of thrombosis and anti-thrombotic therapies. Thrombosis Journal. 2016.

2 Jamasbi J, Ayabe K, Goto S, Nieswandt B, Peter K, Siess W. Platelet receptors as therapeutic targets: Past, present and future. Thromb Haemost 2017; 117: 1249–57.

3 Dunne E, Spring CM, Reheman A, Jin W, Berndt MC, Newman DK, Newman PJ, Ni H, Kenny D. Cadherin 6 has a functional role in platelet aggregation and thrombus formation. Arterioscler Thromb Vasc Biol 2012; 32: 1724–31.

4 Elrod JW, Park JH, Oshima T, Sharp CD, Minagar A, Alexander JS. Expression of junctional proteins in human platelets. Platelets 2003; 14: 247–51.

5 Scanlon VM, Teixeira AM, Tyagi T, Zou S, Zhang PX, Booth CJ, Kowalska MA, Bao J, Hwa J, Hayes V, Marks MS, Poncz M, Krause DS. Epithelial (E)-Cadherin is a Novel Mediator of Platelet Aggregation and Clot Stability. Thromb Haemost Georg Thieme Verlag; 2019; 119: 744–57.

6 Halbleib JM, Nelson WJ. Cadherins in development: cell adhesion, sorting, and tissue morphogenesis. Genes Dev Cold Spring Harbor Laboratory Press; 2006; 20: 3199–214.

7 Sotomayor M, Gaudet R, Corey DP. Sorting Out a Promiscuous Superfamily: Towards Cadherin Connectomics.

8 Bartolomé RA, Peláez-García A, Gomez I, Torres S, Fernandez-Aceñero MJ, Escudero-Paniagua B, Imbaud JI, Casal JI. An RGD Motif Present in Cadherin 17 Induces Integrin Activation and Tumor Growth *. in Press; 2014;.

9 Ruoslahti E, Pierschbacher MD. New perspectives in cell adhesion: RGD and integrins. Science (80-) American Association for the Advancement of Science; 1987; 238: 491–7.

10 de la Fuente M, Noble DN, Verma S, Nieman MT. Mapping human protease-activated receptor 4 (PAR4) homodimer interface to transmembrane helix 4. J Biol Chem 2012; 287: 10414–23.

11 Arachiche A, Mumaw MM, de la Fuente M, Nieman MT. Protease-activated receptor 1 (PAR1) and PAR4 heterodimers are required for PAR1-enhanced cleavage of PAR4 by α-thrombin. J Biol Chem 2013; 288: 32553–62.

12 Arachiche A, de la Fuente M, Nieman MT. Calcium Mobilization And Protein Kinase C Activation Downstream Of Protease Activated Receptor 4 (PAR4) Is Negatively Regulated By PAR3 In Mouse Platelets. PLoS One 2013; 8: e55740.

13 Li W, McIntyre TM, Silversteinc RL. Ferric chloride-induced murine carotid arterial injury: A model of redox pathology. Redox Biol Elsevier B.V.; 2013; 1: 50–5.

14 Li W, Gigante A, Perez-Perez MJ, Yue H, Hirano M, McIntyre TM, Silverstein RL. Thymidine phosphorylase participates in platelet signaling and promotes thrombosis. Circ Res Lippincott Williams and Wilkins; 2014; 115: 997–1006.

15 Mumaw MM, de la Fuente M, Noble DN, Nieman MT. Targeting the anionic region of human protease-activated receptor 4 inhibits platelet aggregation and thrombosis without interfering with hemostasis. J Thromb Haemost 2014; 12: 1331–41.

16 Bouck EG, Zunica ER, Nieman MT. Optimizing the presentation of bleeding and thrombosis data: Responding to censored data using Kaplan-Meier curves. Thromb Res Elsevier; 2017; 158: 154–6.

17 Mège RM, Ishiyama N. Integration of cadherin adhesion and cytoskeleton at adherens junctions. Cold Spring Harbor Perspectives in Biology. Cold Spring Harbor Laboratory Press; 2017. p. a028738.

18 Chimori Y, Hayashi K, Kimura K, Nishida W, Funahashi SI, Miyata S, Shimane M, Matsuzawa Y, Sobue K. Phenotype-dependent expression of cadherin 6B in vascular and visceral smooth muscle cells. FEBS Lett 2000; 469: 67–71.

19 Pippucci T, Maresca A, Magini P, Cenacchi G, Donadio V, Palombo F, Papa V, Incensi A, Gasparre G, Valentino ML, Preziuso C, Pisano A, Ragno M, Liguori R, Giordano C, Tonon C, Lodi R, Parmeggiani A, Carelli V, Seri M. Homozygous NOTCH 3 null mutation and impaired NOTCH 3 signaling in recessive early-onset arteriopathy and cavitating leukoencephalopathy. EMBO Mol Med 2015; 7: 848–58.

20 Haemmerle M, Stone RL, Menter DG, Afshar-Kharghan V, Sood AK. The Platelet Lifeline to Cancer: Challenges and Opportunities. Cancer Cell 2018;.

21 Bialucha CU, Collins SD, Li X, Saxena P, Zhang X, Dürr C, Lafont B, Prieur P, Shim Y, Mosher R, Lee D, Ostrom L, Hu T, Bilic S, Rajlic IL, Capka V, Jiang W, Wagner JP, Elliott GN, Veloso A, et al. Discovery and optimization of HKT288, a cadherin-6-targeting ADC for the treatment of ovarian and renal cancers. Cancer Discov American Association for Cancer Research; 2017; 7: 1030–45.

22 Paul R, Ewing CM, Robinson JC, Marshall FF, Johnson KR, Wheelock MJ, Isaacs WB. Cadherin-6, a cell adhesion molecule specifically expressed in the proximal renal tubule and renal cell carcinoma. Cancer Res 1997; 57: 2741–8.

23 Bartolomé RA, Aizpurua C, Jaén M, Torres S, Calviño E, Imbaud JI, Casal JI. Monoclonal Antibodies Directed against Cadherin RGD Exhibit Therapeutic Activity against Melanoma and Colorectal Cancer Metastasis. Clin Cancer Res American Association for Cancer Research; 2018; 24: 433–44.

